# Experimental Methods for CRISPR Enzyme Assays with Fluorescence Readout

**DOI:** 10.64898/2026.06.03.729647

**Authors:** Qi Jiang, Alexandre S Avaro, Hyunjun Bae, Angelina Sorensen, Juan G Santiago

**Affiliations:** Department of Mechanical Engineering, Stanford University, Stanford, USA; Laboratoire d’Hydrodynamique (LadHyX), CNRS, École Polytechnique, Institut Polytechnique de Paris, 91120 Palaiseau, France; Institut Pasteur, Université Paris Cité, Physical Microfluidics and Bioengineering, 25-28 Rue du Dr. Roux, 75015 Paris, France

**Author notes:** Corresponding author: Juan G. Santiago, Department of Mechanical Engineering, Stanford University, Stanford, CA 94305, Office.

## Abstract

Fluorescence-based CRISPR diagnostic assays have become a popular platform for nucleic acid detection due to their programmability, configurability, specificity, and compatibility with standard laboratory equipment. However, reported enzymatic kinetic rates and limits of detection for CRISPR *trans-*cleavage assays vary by several orders of magnitude across the literature. This variation in performance parameters is coupled with and exacerbated by inconsistent calibration, incomplete correction of measurement biases, and nonstandardized or incomplete data-analysis procedures. We present an experimental protocol and quantitative analysis framework for fluorescence-based enzyme assays using routine laboratory instrumentation, including thermocyclers and fluorescence microplate readers. Building on previous studies of CRISPR enzyme kinetics and fluorescence calibration, we describe procedures for flat-field and background correction; comprehensive fluorescence calibration including correction for inner-filter-effect; quantification and implications of reporter degradation; extraction of Michaelis-Menten kinetic parameters; and determination of assay limits of detection. We provide step-by-step experimental guidelines and open-source Python implementations for each stage of the workflow. Using representative Cas12 *trans-*cleavage datasets, we demonstrate that explicit fluorescence calibration and correction procedures substantially reduce systematic bias in measured kinetic rates and improve consistency between experiments. Our framework aims to establish standardized practices for quantitative fluorescence-based CRISPR assays and provides researchers with practical tools for reproducible kinetic characterization and rational assay design.

## 1. Introduction

Enzymatic assays are a workhorse of biochemistry for the identification of nucleic acid sequences and the quantification of their concentrations. The most influential and prevalent of such assays are enzyme assays which offer exponential amplification of specific nucleic acid sequences. The most important of these is the polymerase chain reaction (PCR) due to its unrivaled sensitivity, strong specificity, reconfigurability, and robustness.^1,2^ Other important exponential amplification enzymatic assays include loop-mediated isothermal amplification (LAMP)^3,4^ and recombinase polymerase amplification (RPA).^5,6^ Linear amplification assays, such as rolling circle amplification (RCA), have also been developed for applications that require low amplification bias.^7^ For all enzyme assays, quantitative, specific, and sensitive implementation requires rigorous experimental procedures for signal calibration, quantification of enzyme kinetics, and rational interpretation of measured parameters. Particularly important is the inference of specific signal from background signal and noise.

Starting with two seminal papers,^8,9^ the last nine years have seen an explosion of the study, development, and implementations of enzymatic assays for detection and identification of nucleic acids based on CRISPR-Cas (clustered regularly interspaced short palindromic repeats, CRISPR-associated).^10–16^ In nearly all CRISPR-based assays, the presence of the target nucleic acid activates a Cas enzyme. Then, in the majority of implementations, the activated Cas is then detected by including synthetic nucleic acids functionalized with a fluorophore and a quencher^8,9^ The type and concentration of these reporter molecules must be judiciously chosen to stoichiometrically favor catalytic cleavage by the activated Cas, while avoiding overly high reporter concentration, which adds non-specific background signal. The cleavage results in a measurable increase in the fluorescence signal. Quantification of the fluorescence intensity value and its rate of increase can each inform on the type and quantity of target molecules in the sample. The sensitivity of all CRISPR-Cas enzyme assays is fundamentally limited by the kinetic rates of the enzyme system,^17^ and so understanding and quantification of kinetic rates is essential.

Despite its great importance and wide applicability, many of the reported experimental methods for CRISPR-Cas nucleic acid assays have been flawed and experimental data have been incomplete and/or lacking self-consistency.^13,18,19^ There have been two main issues. First, initial quantifications of enzyme kinetic rates for CRISPR-Cas have demonstrable errors and/or a lack of self-consistency.^13,18,19^ The discrepancy cannot be accounted for by experimental uncertainty.^20^ For example, reports of the catalytic conversion rate by CRISPR-Cas 12 and 13 vary by over 8 orders of magnitude and have been demonstrably inconsistent by factors of 1,000 and higher. Second, and related, many of the limits of detection that have been reported for CRISPR-Cas assays are very difficult to reconcile with the observed low kinetic rates of CRISPR-Cas.^13,19^ This is particularly (and nearly exclusively) an issue with CRISPR-Cas assays which attempt to measure and identify trace analytes without the help of “pre-amplification” of nucleic acid targets with traditional enzyme amplification techniques like PCR or LAMP. Recently, Avaro and Santiago^13^ reviewed CRISPR-Cas assays, kinetic rate measurements, limits of detection, and the possible sources of discrepancies. Drawing on scaling, back-of-the-envelope estimates, and published corrections, they identified fluorescence signal calibration as the most common source of discrepancy in kinetic rate determination. The latter work also reviewed experimental considerations for fluorescence-based CRISPR enzyme assays, including development of calibration methods, and provided recommendations for quantitative reporting and interpretation of CRISPR-Cas assays.

In this work, we aim to provide standard experimental methods and protocols for CRISPR-based assays that use fluorescence readout. We report a fairly exhaustive protocol for signal calibration which accounts for the signal of uncleaved reporters and includes both the inner-filter effect and flat-field corrections. We provide detailed procedures to quantify enzymatic kinetic rates and to experimentally determine the diagnostic assay limit of detection. Our discussions are focused on efforts using routine laboratory equipment. We also provide open-source Python codes to extract figures of merit from raw fluorescence data. Together, these resources provide a framework which may improve the accuracy and reproducibility of CRISPR-based assay characterization.

## 2. Protocol overview

The current protocol paper begins with a summary of the Michaelis-Menten kinetic framework in the context of the *trans*-cleavage reaction of CRISPR assays (Section 3). We then provide recommendations on reagents and equipment (Section 4). In Sections 5 and 6, we describe calibration methods which are essential for any study. As shown in **Figure 1**, the discussions and content then separate into two parallel tracks where we discuss two main types of CRISPR assays. First (top row of **Figure 1**), we describe the determination of kinetic rate parameters. Second (bottom row), we discuss improvement and determination of an assay limits of detection (LoD). To measure Michaelis-Menten kinetic rates (Section 7), CRISPR reactions are carried out using varying substrate concentrations to extract the rate constants. The consistency between the experimental procedure and the resulting rates is verified using three simple back-of-the-envelope calculations (Section 8). For the determination of limits of detection assay (Section 9), CRISPR reactions are performed using varying activator concentrations. Finally, we briefly comment on the performance assessment of CRISPR assays conducted within miniaturized compartments (Section 10).

**Figure 1.**
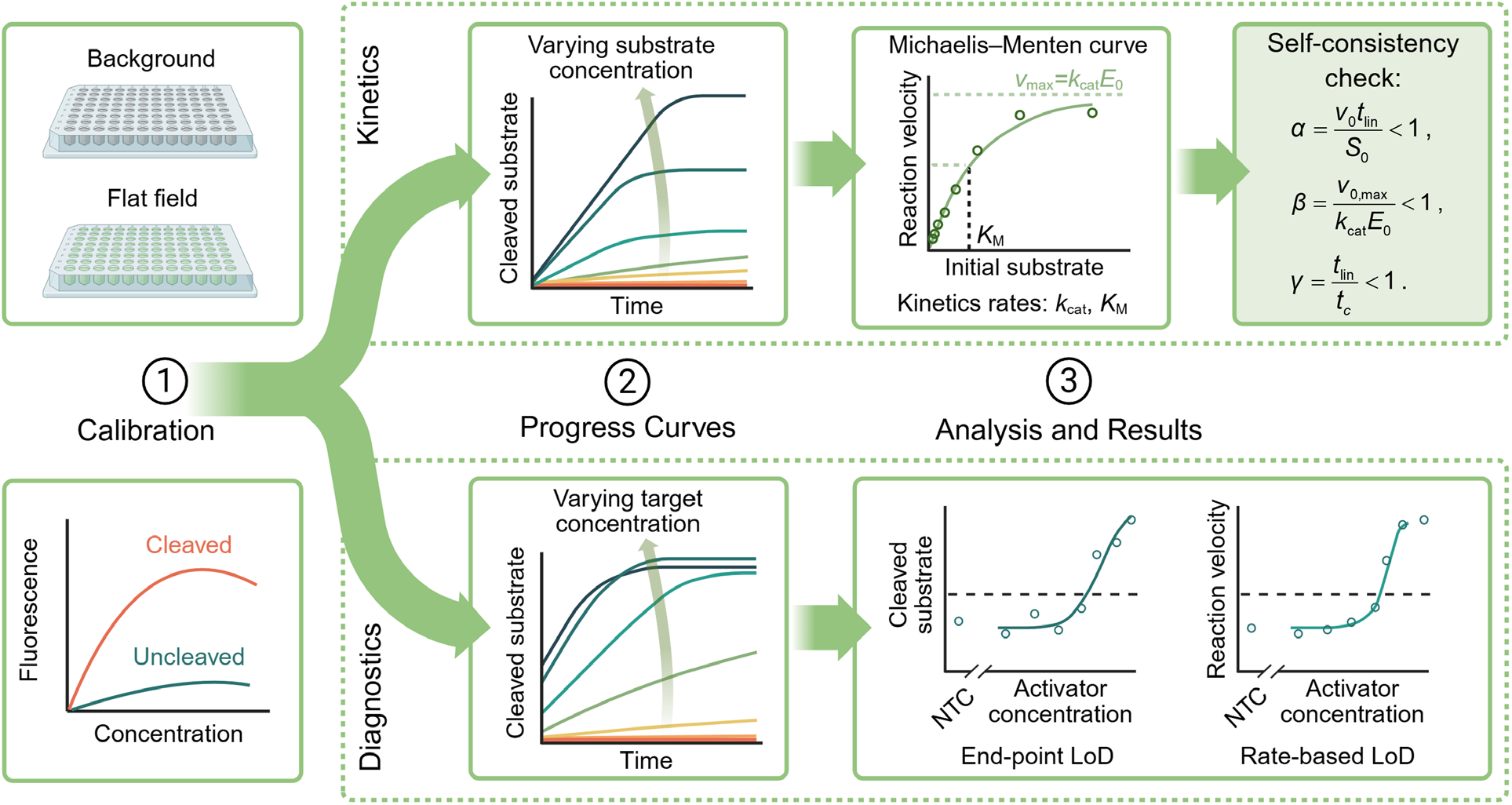
Workflow diagram for two types of enzymatic assays with fluorescence-based readout using a thermocycler. The first assay (top row) describes measurement of kinetic rates, and the second (bottom row) is a limit-of-detection measurement. Both pipelines begin with flat-field and background correction of raw fluorescence images produced by the thermocycler array (Section 5). This is followed by fluorescence signal calibration, to convert corrected fluorescence signal to molar concentrations of cleaved versus uncleaved reporters, and to extract the reporter degradation rate *k*_rep_ (Section 6). The workflow then branches. For the quantification of enzyme kinetics, calibrated progress curves at varying initial substrate concentrations are obtained and compared to the Michaelis-Menten model to extract the catalytic turnover rate *k*_cat_ and the Michaelis constant *K*_M_ (Section 7). Resulting kinetic parameters should be checked for self-consistency using back-of-the-envelope parameters *α, β*, and *γ* (Section 8). Conversely, to assess the limit of detection of an assay, calibrated progress curves at varying initial target concentrations are obtained and processed to quantify limits of detection (LoDs) using either endpoint fluorescence values or initial reaction velocities (Section 9).

## 3. Kinetic model: Michaelis-Menten framework for CRISPR *trans*-cleavage

Fluorescence-based CRISPR assays typically involve two sequential processes: *cis-*cleavage and *trans-*cleavage. In the *cis-*cleavage step, a target nucleic acid binds to the CRISPR-associated enzyme through guide RNA (gRNA) recognition and the target nucleic acid is cleaved by the enzyme. This reaction activates the enzyme. Following activation, the CRISPR ribonucleoprotein (RNP) undergoes *trans-*cleavage activity where it indiscriminately cleaves nucleic acids. The stoichiometry of fluorophore-quencher reporter molecules is designed so that the cleaving of reporters dominates the *trans-*cleavage rates. Hence, reporter molecules are the substrate of the enzyme. Cleavage of reporters creates a physical separation between the fluorophore (on one end of the reporter) and quencher (on the other end). The cleavage relieves quenching and restores the fluorophore’s intrinsic quantum yield.^21^ This produces an increase in fluorescence signal that can be quantified as a measure of the concentration of cleaved reporter molecules (i.e., the reaction product). The *trans-*cleavage process is a multi-turnover enzymatic reaction and is nearly always the rate-limiting step governing signal generation.^18^

The Michaelis-Menten enzyme kinetics framework is, to date, the most successful model for the rate-limiting *trans-*cleaveage kinetics of CRISPR-Cas 12 and Cas 13 enzymes.^22–24^ The model involves an enzyme *E*, uncleaved substrate *S*, enzyme-substrate intermediate complex *C*, and cleaved reporter product *P*:

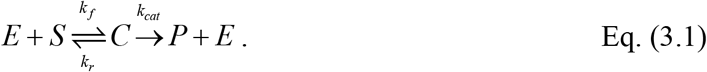

Here, *k*_*f*_ and *k*_*r*_ are the forward and reverse rate constants for enzyme-substrate binding, and *k*_cat_ is the catalytic turnover rate.^18,22^ In CRISPR assays, *E* (“enzyme”) represents the activated Cas-gRNA complex, *S* (“substrate”) is the uncleaved fluorophore-quencher reporter, and *P* (“product”) is the cleaved reporter molecule. As mentioned above, cleavage of reporters separates fluorophore and quencher, producing an increase in fluorescence signal. The corresponding system of reaction-rate equations is:

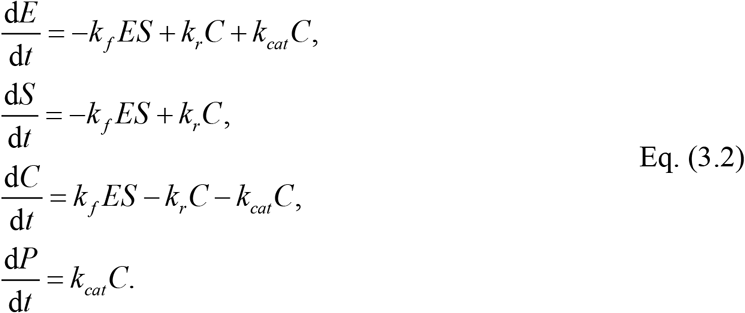

The initial conditions for CRISPR *trans*-cleavage experiments are:

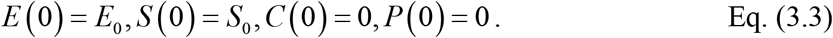

Under the quasi-steady-state and reactant-stationary assumptions,^25^ the reaction velocity can be approximated by the classical Michaelis-Menten expression

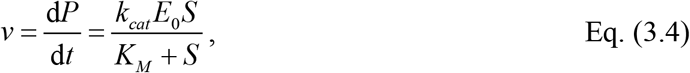

where the Michaelis constant is defined as *K*_*M*_ = (*k*_*cat*_ + *k*_*r*_) / *k* _*f*_.

In this framework, *k*_cat_ is the turnover number and governs the maximum catalytic turnover rate of the activated enzyme, whereas *K*_M_ represents the reporter concentration at which the reaction velocity reaches half of its maximum value. The ratio *k*_cat_ / *K*_M_ is commonly referred to as catalytic efficiency. The initial concentration of activated enzyme *E*_0_ is approximately equal to the concentration of target molecules present in the reaction. As described by Huyke et al.,^17^ and Avaro and Santiago,^20^ *k*_cat_ / *K*_M_ (and not *k*_cat_) very often governs the sensitivity of CRISPR-Cas enzyme assays. This is due to a trade-off between *trans*-cleavage reaction production rate and background signal increase which we shall discuss below in Section 9. A well-designed CRISPR-Cas detection assay should use a reporter concentration that is much higher than the target concentration, but overly high reporter concentration decreases signal-to-background ratio and lowers limit of detection.

In the following sections, we build on this framework to describe procedures for fluorescence calibration, extraction of progress curves, estimation of kinetic parameters, and determination of assay limits of detection from fluorescence-based CRISPR experiments.

## 4. Material Selection and Equipment

We here summarize common reagents, instruments, and other practical considerations for fluorescence-based CRISPR assays. We focus on materials and equipment compatible with quantitative fluorescence measurements and fluorescence calibration. A more comprehensive list of recommended reagents and equipment is provided in Section S1 in the Supporting Information (SI).

Oligonucleotides may be ordered either lyophilized or diluted in nuclease-free water. We recommend quantifying nucleic acid concentrations using a spectrophotometer (e.g., NanoDrop, ThermoFisher Scientific) or a fluorometer (e.g., Qubit, ThermoFisher Scientific). This is particularly important for reactions in which nucleic acids are limiting reagents. gRNAs are susceptible to nuclease degradation and repeated freeze-thaw cycles. We therefore recommend aliquoting gRNA stocks upon receipt and storing aliquots at −80°C to minimize freeze-thaw cycles. Target DNA and fluorophore-quencher reporters may be aliquoted and stored at −20°C. Fluorophore-quencher molecules should be protected from light. We recommend avoiding large excesses of gRNA relative to Cas protein, particularly for assays using ssDNA activators. The reason is that excess free gRNA may hybridize with target ssDNA molecules and reduce the fraction of target available for Cas activation. In previous work, Jiang et al.^26^ showed a [gRNA]:[Cas] ratio between 0.5 and 2.0 produced minor changes in measured Cas12 *trans*-cleavage rates. Nevertheless, we recommend avoiding [gRNA]:[Cas] ratios higher than 1 to minimize potential hybridization of free gRNA with target ssDNA. We recommend mixing the reagents on ice, particularly for fast reactions where mixing time is not negligible compared to the reaction timescale.^22^ This reduces reporters cleaved prior to measurement of reaction rates. We provide specific guidelines and back-of-the-envelope estimates to avoid such a bias in Section 9 below.

We next discuss fluorescence detection platforms used for quantitative CRISPR enzyme assays and fluorescence calibration. The current authors have typically used qPCR or plate reader platforms for real-time kinetic measurements. Such platforms support extended isothermal holds at 37 °C, thermal uniformity across wells within ±0.5 °C, and heated lids that reduce evaporation over the typical 10 to 120 min assays. The most consequential distinction among such instruments is not the vendor but the optical architecture. Whole-plate camera systems (e.g., Applied Biosystems QuantStudio and 7500 Fast, Roche LightCycler 480 II) use a stationary CMOS or CCD detector with integrated uniformity calibration and passive reference normalization to correct for optical vignetting. This reduces reliance on a passive reference dye (e.g., ROX) for well-to-well normalization.^27^ In our experience, however, neither architecture eliminates nonuniformity of measured signals across wells.^13,28^ We therefore recommend the flat field and background corrections described in Section 4 on every platform. Some modern instruments include factory calibration and passive reference normalization, such as the QuantStudio platforms (Applied Biosystems, Thermo Fisher, 2022).^29^ These integrated corrections help reduce but do not eliminate signal variation, and so we nevertheless recommend the process in Section 5.

Lastly, we offer some comments around assays performed using systems other than typical thermal cyclers, such as custom imaging devices or microfluidic devices. We distinguish among three classes which are commonly used for CRISPR diagnostics. First, scientific CMOS (sCMOS) cameras provide spatially resolved readouts on microscopy platforms, and these are useful for microfluidic-based assays. The latter include, among others, droplet and digital CRISPR assays. We recommend a sensor with a quantum efficiency above 70% and read noise below 2 e^−^ rms. For instance, our laboratory has used a Hamamatsu ORCA-Flash 4.0 sCMOS to quantify CRISPR reaction rates in on-chip isotachophoresis.^30,31^ Ackerman et al. used the same camera in their CARMEN-Cas13 platform.^32^ Second, photodiodes provide an inexpensive and single channel readout with minimal optical complexity at the cost of spatial information. A typical setup pairs the photodiode with an LED source, an optical emission filter, and a transimpedance amplifier.^33,34^ Third, smartphone camera CRISPR readers have been reported. Current implementations often apply little optical calibration.^14^ For example, reported assays have used various image processing approaches, but without flat-field or background correction.^35–37^ Regardless of detector class, we recommend consideration of the techniques described in Sections 5 and 6.

## 5. Flat-field and Background Correction

The measured fluorescence signal across microwells used in real-time thermocyclers and plate readers is not uniform across the microwell array even for uniform fluorophore concentrations. These differences may be due to non-uniformities in illumination, light-capture optics, and detector response, among other factors. This issue is exacerbated by non-uniformities in the dispensing of liquids (e.g., pipetting errors). In practice, the current authors find that even careful filling of the same volume of the same solution across the array can result in up to 30% differences in the measured signal.^13^ Further, a substantial amount of the measurable nonuniformity is repeatable across experiments over periods of one to two weeks. Interestingly, we have observed such non-uniformity of response even immediately after servicing of the instrument by the manufacturer.^13^ We therefore here suggest a correction for the repeatable non-uniformity of measured fluorescence signal.

The correction is analogous to the flat-field correction that is often used in imaging and particularly in fluorescence microscopy.^20,28,38^ We recommend applying this correction prior to any fluorescence-to-concentration conversion (see Section 6).

The corrected signal in each well at each time point is

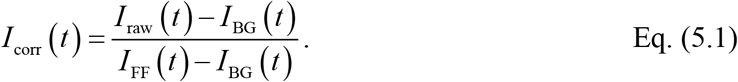

Here, each *I*_*i*_ (*t*) represents a time-dependent array corresponding to the microwells of the instrument. *I*_raw_ is the raw fluorescence signal from the experiment, *I*_BG_ is the background signal acquired with reaction buffer alone (no fluorescent probes), *I*_FF_ is the flat-field signal acquired with a uniform fluorophore solution, and *I*_corr_ is the corrected signal. The quantities *I*_raw_, *I*_BG_, and *I*_FF_ each have units of relative fluorescence units (RFU), whereas *I*_corr_ is dimensionless. All data should be collected within the same instrument at the same temperature of interest and *I*_FF_ should be taken at reagent concentrations on the same order as those used in the kinetic experiment. The correction defined by Eq. (5.1) is applied at every time point using the corresponding arrays *I*_FF_ (*t*) and *I*_BG_ (*t*). Therefore, *I*_FF_ (*t*) and *I*_BG_ (*t*) should be acquired using the same instrument settings and sampling interval as *I*_raw_ (*t*). The intent of this correction is to normalize the measured response in terms of the background as the “noise floor” and to correct for the non-uniformity in signal contained within array *I*_FF_ (*t*). As an example, **Figures 2A-2C** display only a single representative frame at *t* = 2142 s for visualization, whereas the correction is applied at every time point using the full time-series arrays *I*_FF_ (*t*) and *I*_BG_ (*t*).

**Figure 2.**
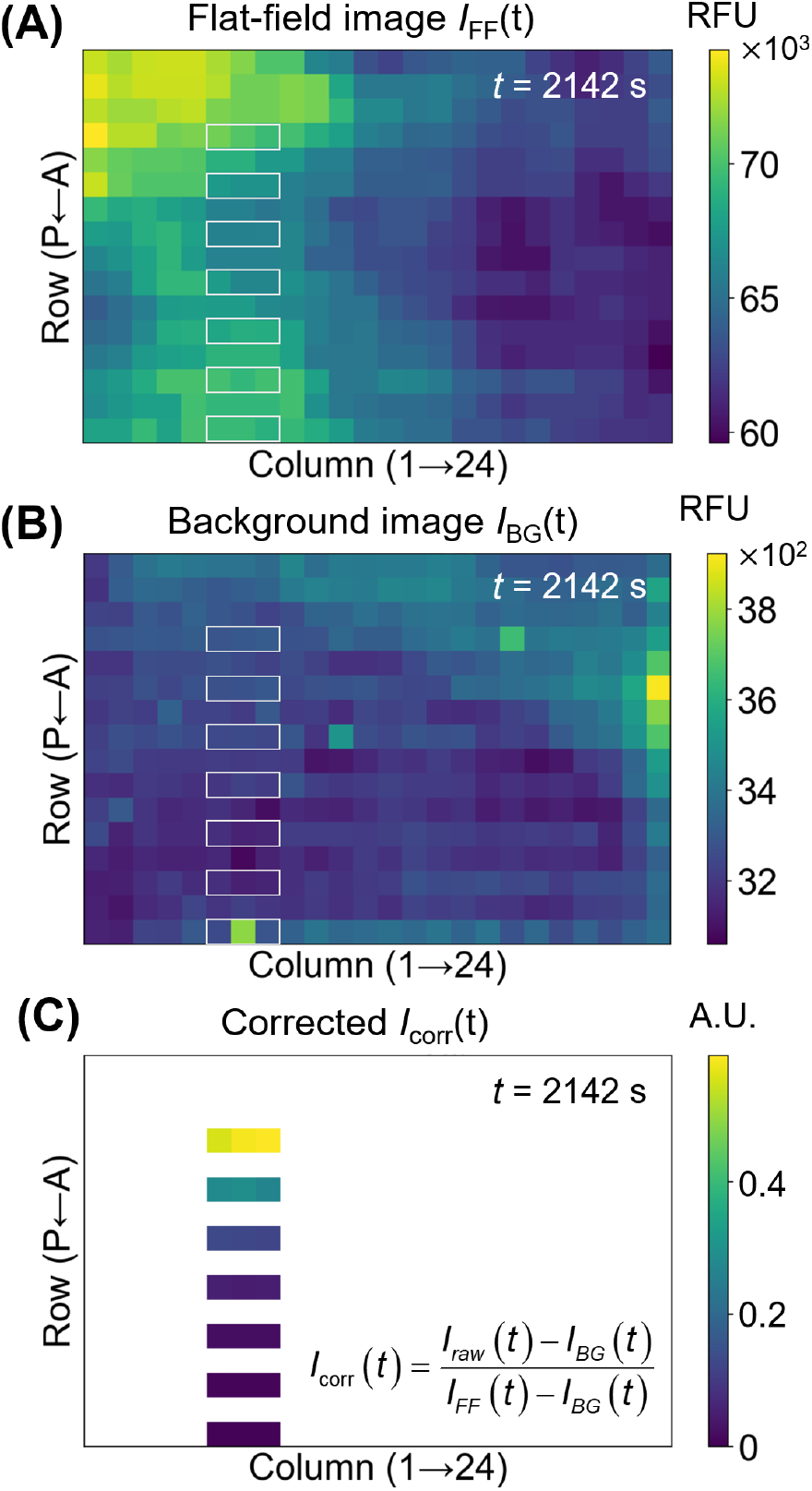
Example experimental flat-field and background correction fluorescence images acquired from a 384-well plate at *t* = 2142 s (cycle 50). **(A)** Flat-field image *I*_FF_.^13^ White rectangles highlight 21 wells used to obtain kinetics data (and so associated with *I*_raw_). **(B)** Background image *I*_BG_. **(C)** Corrected signal *I*_corr_ calculated using Eq. (5.1) for the highlighted wells. The kinetic experiment used 21 wells arranged as seven triplicates, with each row corresponding to a single reporter concentration.

**Figure 2A** shows a flat-field image *I*_FF_ acquired on a 384-well plate where each well contains 20 of 10 μM fluorescein solution.^13^ The flat-field images can also be generated using the cleaved reporter concentration required for calibration (see Section 6). For *I*_FF_, we sealed the wells as uniformly as possible using a single optical film. In these data, the flat field signal varies from 7.4 × 10^4^ RFU in the upper-left wells to 6.0 × 10^4^ RFU in the lower-right wells, a relative spread of 23%. The white rectangles highlight the wells whose data we track through the correction.

**Figure 2B** shows the corresponding background image *I*_BG_ at the same time point. We acquired *I*_BG_ using the same volume of nuclease-free water. *I*_BG_ ranges from 3.1 × 10^3^ to 3.8 × 10^3^ RFU (20% variation) and interestingly shows a few isolated hot spots that do not align with the spatial pattern of *I*_FF_. We hypothesize these features arise from spatially localized autofluorescence of the optical film and/or well array plastic.

**Figure 2C** shows the corrected signal *I*_corr_ at *t* = 2142s using equation (5.1) for the highlighted region. We used 21 wells in the kinetic experiment arranging as seven rows of triplicates, where each row contained three wells dispensed with the same reporter concentration. The remaining wells of the plate were not part of this measurement and were left blank. After correction, the coefficient of variation (CV) across triplicates decreased for 6 of 7 concentrations (Figure S1), consistent with reduction of position-dependent systematic bias from the measured signal. We hypothesize the residual variability reflects random pipetting errors and detector non-uniformity. Note that the flat-field and background signals are not constant in time (See Figure S1). We attribute this temporal variation primarily to slow photobleaching of the fluorescent dye which was used to acquire *I*_FF_.

## 6. Fluorescence Signal Calibration

After correction for (repeatable) non-uniform flat-field and background, the corrected signal *I*_corr_ is dimensionless. To convert *I*_corr_ to molar concentrations of cleaved reporter *P*, an experimentalist needs a relation between measured fluorescence intensities and species concentration. This calibration model should capture the fluorescence of both cleaved and uncleaved reporters. Note that uncleaved reporters have measurable and significant fluorescence, but with a lower quantum yield than cleaved reporters. The calibration should also take into account any significant inner-filter effect (IFE). The IFE arises from self-absorption of excitation and emission photons by all molecules in the solution and produces a non-monotonic relationship between *I*_corr_ and concentration.^28,39^ The IFE is particularly important to calibration of CRISPR-Cas assays with fluorescent readout because of the use of quencher groups. IFE is strongly affected by the presence of quencher molecules of both uncleaved and cleaved reporters, and the effect is severe at high reporter concentrations. In the following sub-sections, we suggest a calibration model which results in quantities required by the rest of the protocol. We first describe an IFE-corrected calibration of cleaved and uncleaved reporter concentrations (Section 6.1). We then quantify a reporter degradation rate *k*_rep_ (Section 6.2).^22^ *k*_rep_ is critical for estimates of the limit-of-detection of the system and therefore for assay design (Section 9). Typically, *k*_rep_ is not needed for accurate determination of Michaelis-Menten kinetic parameters *k*_cat_ and *K*_M_ (Section 7).

### 6.1 Calibration of cleaved and uncleaved reporters

Avaro and Santiago^13^ reviewed the history of calibration methods for CRISPR-based assays. A general calibration model describes wells containing mixtures of cleaved and uncleaved reporters is the following:

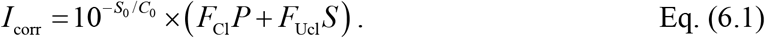

Here *F*_Cl_ and *F*_Ucl_ are the per-concentration calibration constants for cleaved and uncleaved reporters, respectively. The factor *C*_0_ is a characteristic concentration that quantifies the strength of the IFE. *C*_0_ is inversely proportional to the product of the molar extinction coefficient and the optical path length, as per the Beer-Lambert law. *S*_0_ denotes the initial reporter concentration. The total reporter concentration in the well is *S*_0_ = *S* + *P*, which is conserved during *trans*-cleavage.

To determine *F*_Cl_, *F*_Ucl_, and *C*_0_, we perform calibration experiments using two dilution series. One series contains fully cleaved reporter at concentration *c*_Cl_, and the other contains fully uncleaved reporter at concentration *c*_Ucl_. In the cleaved series, *S* = 0 and *P* = *c*_Cl_. In the uncleaved series, *P* = 0 and *S* = *c*_Ucl_. Eq. (6.1) therefore reduces to

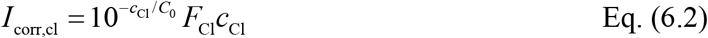

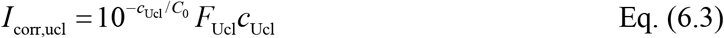

We then jointly fit both datasets using nonlinear least squares to determine the three calibration parameters *F*_Cl_, *F*_Ucl_, and the shared parameter *C*_0_. Once *F*_Cl_ and *F*_Ucl_ are obtained, the conversion of fluorescence values to molar concentration of cleaved reporters followed the equation:

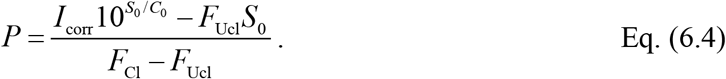

**Figure 3A** shows a representative calibration of corrected fluorescence as a function of reporter concentration for cleaved (green) and uncleaved (red) reporters.^13^ The corrected fluorescence increases monotonically with reporter concentration for both reporters within the range. However, the fluorescence signals level off at higher concentration as IFE attenuates the signal. Dashed lines show the linear extrapolation from low concentration, while solid lines show the IFE-corrected nonlinear fit to Eqs. (6.2) and (6.3). The deviation between the dashed and solid green curves quantifies the IFE attenuation of the cleaved-reporter signal. The fit yields *C*_0_ = 4.7 μM. *C*_0_ is typically on the order of a few micromolar for short fluorophore-quencher reporters in standard CRISPR buffers.

**Figure 3.**
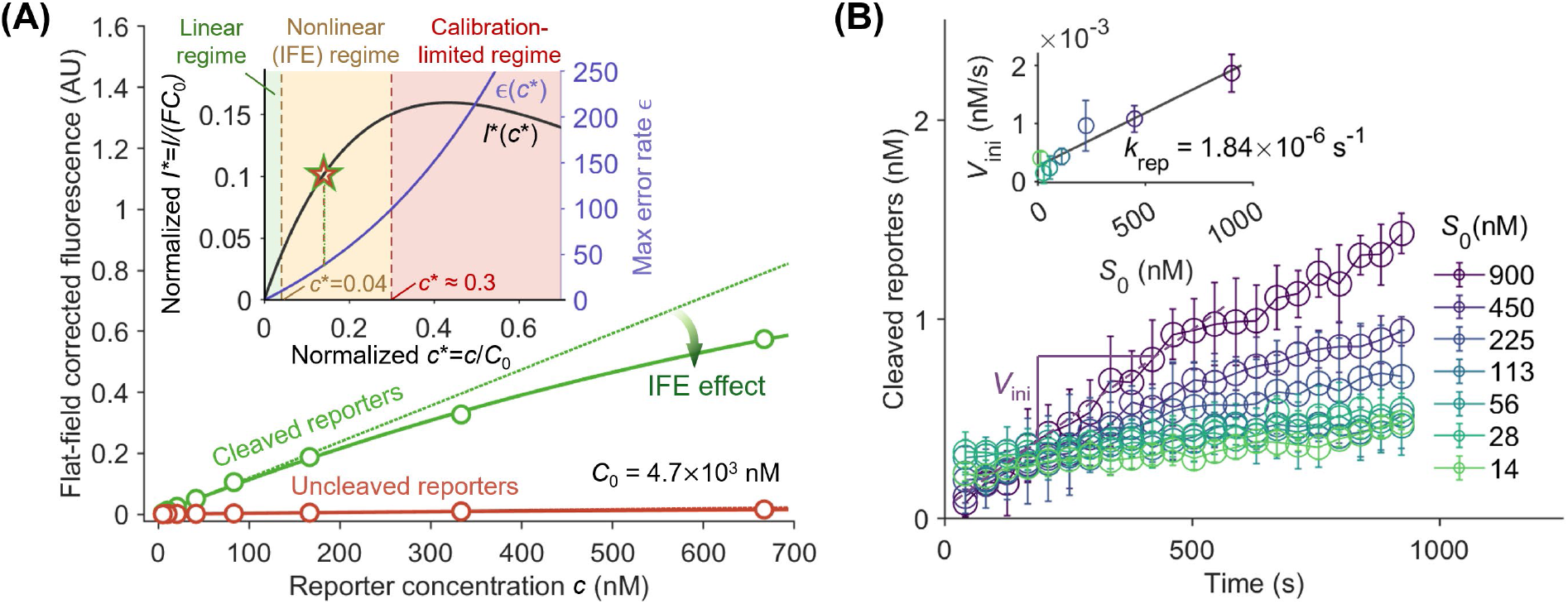
Calibration relating corrected fluorescence to reporter concentration and quantification of the reporter degradation rate. **(A)** Flat-field corrected fluorescence as a function of reporter concentration for cleaved (green) and uncleaved (red) reporters.^13^ Symbols refer to experimental measurements. Solid lines are nonlinear least-squares fits to Eqs. (6.2) and (6.3) (which accounts for IFE). Dotted lines are the linear extrapolation from low concentration. The dark green arrow highlights the deviation between the IFE-corrected fit and the linear extrapolation. The inset shows the normalized signal *I*^*^ (black, left ordinate) and the associated maximum error rate *ε* incurred by neglecting IFE (purple, right ordinate), each plotted against the normalized concentration *c*^*^. The non-monotonic shape of *I* ^*^ (*c*^*^)defines three regimes: a useful linear regime (*c*^*^ < 0.04, green), a nonlinear IFE regime (0.04 < *c*^*^ < 0.3, orange), and calibration-limited regime (*c*^*^ > 0.3). *I*^*^ reaches a peak at *c*^*^ = 0.43 (not shown). **(B)** Cleaved reporter concentration as a function of time for seven values of the initial reporter concentration *S*_0_ ranging from 14 to 900 nM, in absence of CRISPR enzyme, gRNA, and target.^22^ The *V* _ini_ noted illustrates the slope used to characterize first-order reporter degradation (Eq. 6.6). The inset plots *V*_ini_ against *S*_0_ across all seven concentrations. Linear fit through the origin yields a degradation rate *k*_rep_ = 1.84 × 10^−6^ s^−1^.

To guide the selection of substrate concentrations for the kinetic assay, we nondimensionalize the calibration curve. The inset of **Figure 3A** shows two curves plotted against the normalized concentration *c*^*^ = *c* / *C*_0_. After normalization, all IFE-corrected calibration curves, including those for cleaved and uncleaved reporters and for various reporter types, collapse onto the same universal curve shown in the inset of **Figure 3A**. The second curve is the maximum error rate *ε* incurred by neglecting the nonlinear effect (purple, right ordinate). The non-monotonic shape of *I* * (*c*\*) defines three operating regimes. In the linear regime (*c*^*^ < 0.04, green shading), *I*^*^ is approximately constant and fluorescence is proportional to concentration. Here the maximum error from neglecting the IFE is below 10%. In the nonlinear IFE regime (0.04 < *c*^*^ < 0.3, orange shading), the relationship between *I*^*^ and *c*^*^ remains single-valued and invertible. Although the calibration curve deviates from linearity due to IFE, Eq. (6.1) can still recover the correct concentration. In the calibration-limited regime (*c*^*^ greater than approximately 0.3, red shading), the calibration curve begins to plateau, and the fluorescence signal becomes insensitive to concentration. At higher *c*^*^, a peak of *I*^*^ occurs at *c*^*^ = 0.43, and the inference of concentration given intensity becomes double-valued. That is, beyond this point, a single value of *I*^*^ corresponds to two possible concentrations, and the calibration can no longer be inverted unambiguously. An experimentalist should avoid this calibration-limited regime when designing kinetic or diagnostic assays. We recommend design assays such that the highest reporter concentration remains below about 0.3 *C*_0_ to avoid the region where the *I*^*^ becomes insensitive to *c*^*^. The star on the *I* * (*c*\*) curve marks the concentration range used in the main plot of **Figure 3A**. It shows that *I* ^*^ (*c*^*^) has not yet reached its peak and the corresponding maximum error rate is 30%.

### 6.2 Quantification of the reporter degradation rate *k*_rep_

Fluorophore-quencher reporters may undergo slow spontaneous degradation in solutions even in the absence of CRISPR enzyme.^22,40^ These processes produce a finite background reaction velocity. This degradation imposes a fundamental floor on the limit of detection of CRISPR assays and must be quantified for any diagnostic application aimed at trace analyte detection (Section 9).

We model reporter degradation in the absence of activated enzyme as a first-order process,

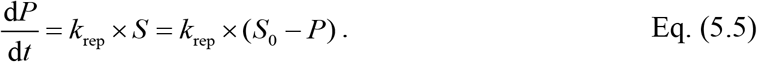

where *k*_rep_ is the first-order degradation rate and *S*_0_ is the initial reporter concentration. For times *t* ≪ 1/ *k*_rep_, the depletion of uncleaved reporter is negligible and the initial reaction velocity simplifies to

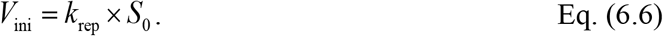

We apply Eq. (6.6) by measuring *V*_ini_ at several values of *S*_0_. We then extract *k*_rep_ from a linear fit.

**Figure 3B** shows an example of reporter-only progress curves measured at seven initial reporter concentration *S*_0_ ranging from 14 to 900 nM.^22^ The experiments were performed without CRISPR enzyme, gRNA, or target. All curves increase approximately linearly over 15 min, consistent with the first-order degradation model described in Eq. (6.6). The annotation *V*_ini_ illustrates the initial-velocity slope for one of the curves. The inset plots *V*_ini_ against *S*_0_ across all seven concentrations and shows a linear dependence. The linear fit yields *k* _rep_ = 1.84 ×10^−6^ s^−1^. Note that *k* _rep_ depends on the reporter chemistry, buffer composition, incubation temperature, and must be remeasured whenever any of these changes.

#### Step-by-step protocol

##### Prepare the cleaved reporter stock

1. Generate fully cleaved reporters using one of two routes. *Option 1*, CRISPR ribonucleoprotein. Incubate the uncleaved reporters at the assigned concentration with activated CRISPR ribonucleoprotein, typically at 20 nM activator, for several hours. *Option 2*, commercial ssDNA endonuclease. Incubate the uncleaved reporters with a non-specific nuclease such as DNase I until the signal saturates.
2. Verify that the fluorescence signal has saturated before use. Acquire fluorescence readings at intervals over the course of the cleavage reaction and confirm that successive readings no longer increase beyond experimental noise.

##### Acquire the flat-field image *I*_FF_

1. Prepare a stock solution of a fluorescent species with excitation and emission wavelengths similar to that of reporters. A good choice for this is cleaved reporter solution (see “prepare the cleaved reporter stock” in Section 6). This should be performed using a concentration that yields a strong but unsaturated signal on the detector.
2. Dispense an identical volume of this solution into every well of the plate.
3. Seal the plate with the same optical film used in subsequent kinetic experiments, briefly centrifuge to remove bubbles, and load the plate into the detector.
4. Acquire fluorescence readings using identical instrument settings (excitation/emission filters, gain, exposure) and the same sampling interval as the kinetic experiment. Acquire *I*_FF_ at every time point sampled in the kinetic experiment to enable a time-resolved correction.

##### Acquire the background image *I*_BG_

1. Prepare 1× reaction buffer without fluorophore, quencher, enzyme, gRNA, or target.
2. Dispense the same volume per well as used for *I*_FF_ and the kinetic experiment.
3. Acquire fluorescence readings using the same instrument settings and sampling interval as *I*_FF_. Acquire *I*_BG_ at every time point sampled in the kinetic experiment.

##### Acquire the raw image *I*_raw_

1. Prepare the experimental plate. *I*_raw_ refers to any raw fluorescence measurement to which the flat-field and background correction will be applied. This includes the reporter calibration data with cleaved and uncleaved reporter dilutions (Section 6.1), the reporter degradation data with reporter dilutions in buffer alone (Section 6.2), the CRISPR reaction kinetics data (Section 7), and the limit-of-detection data with titrated activator concentrations (Section 9).
2. Dispense the same volume per well as used for *I*_FF_ and *I*_BG_.
3. Seal the plate, briefly centrifuge to remove bubbles, and load the plate into the detector.
4. Acquire fluorescence readings using the same instrument settings and sampling interval as *I*_FF_ and *I*_BG_.

##### Calculate the corrected image *I*_corr_

1. Apply Eq. (5.1) elementwise across all wells and all time points to obtain *I*_corr_ (*t*) from *I*_raw_ (*t*), *I*_BG_ (*t*), and *I*_FF_ (*t*).
2. Verify that *I*_FF_ (*t*), *I*_BG_ (*t*), and *I*_raw_ (*t*) are sampled on the same temporal grid before applying the correction.
3. We provide an open-source implementation^41^ of this calculation: see **Script 1** (c.f. Section S4 in SI) for the flat-field and background correction routine, which takes *I*_FF_ (*t*), *I*_BG_ (*t*) and *I*_raw_ (*t*) as input and returns *I*_corr_ (*t*).

##### Acquire the calibration data for both cleaved and uncleaved reporter solutions

1. Choose a range of reporter concentrations so as the highest concentration spans well into the IFE regime and the calibration captures *C*_0_ reliably; we typically use a highest concentration of 1 to 5 μM using 7 to 8 dilution points.
2. Prepare a parallel 2-fold serial dilution series of the fully uncleaved reporter in 1× reaction buffer over the same concentration range.
3. Dispense each dilution into triplicate wells of the same plate, using the same volume per well as the intended kinetic experiment.
4. Seal the plate with the same optical film, briefly centrifuge to remove bubbles, and load the plate into the detector.
5. Acquire *I*_raw_ using the same instrument settings and sampling interval as the kinetic experiment (Section 5). Acquire *I*_FF_ and *I*_BG_.

##### Compute the calibration parameters

1. Apply the flat-field and background correction (Eq. 5.1) to obtain *I*_corr_ for each calibration well at each time point.
2. Average *I*_corr_ across the time points of the calibration acquisition for each well to suppress per-frame noise. The corrected signal of cleaved and uncleaved reporters in 1× buffer should be approximately constant over the calibration acquisition.
3. Fit Eqs. (6.2) and (6.3) jointly to the cleaved and uncleaved calibration data using nonlinear least squares to extract *F* _Cl_, *F* _Ucl_, and *C*_0_. We provide an open-source implementation^41^ of this calibration: see **Script 2** in SI for the IFE-aware joint fit, which takes the cleaved and uncleaved calibration data as input and returns *F*_Cl_, *F*_Ucl_, and *C*_0_.
4. Verify that the fitted *C*_0_ is physically reasonable, on the order of a few μM for typical short fluorophore-quencher reporters. Compute 0.3 *C*_0_ to obtain the maximum reporter concentration usable in subsequent kinetic and diagnostic experiments.

##### Acquire the reporter-degradation data (for Diagnostics Purpose)

1. Prepare a serial dilution of intact uncleaved reporter in 1× reaction buffer, omitting the CRISPR enzyme, gRNA, and target. Could use the same sets of concentration as the kinetic experiment.
2. Dispense each dilution into triplicate wells on the same plate, using the same volume per well as the kinetic experiment.
3. Seal the plate with the same optical film, briefly centrifuge to remove bubbles, and load the plate into the detector.
4. Incubate at the kinetic-experiment temperature (typically 37 °C) and acquire *I*_raw_ using the same instrument settings and sampling interval as the kinetic experiment. Sample for 15 min.
5. Acquire *I*_FF_ and *I*_BG_.

##### Compute *k*_rep_ (for Diagnostics Purpose)

1. Apply the flat-field and background correction (Eq. 5.1) and the IFE-aware calibration (Eq. 5.4) to convert *I*_raw_ at each well to cleaved reporter concentration *P*(t).
2. For each *S*_0_, fit *P*(t) using linear regression and record the slope as *V*_ini_.
3. Plot *V*_ini_ against *S*_0_ across all dilutions and perform a linear fit. The slope is *k*_rep_, with units of s^−1^. See **Script 2**^41^ in SI for open-source script.

## 7 Measurement of kinetic rates

We here focus on the quantification of *trans*-cleavage kinetics. We assume the enzyme kinetics follow the Michaelis-Menten model introduced in Section 3. With the flat-field and background signal arrays and the calibration parameters, the raw fluorescence signal can then be converted into calibrated cleaved-reporter progress curves *P*(*t*) in molar units. We proceed to extract estimates of kinetic rate parameters from the experimental data.

*Trans*-cleavage progress curves are measured at varying initial substrate (of the uncleaved reporter) concentrations *S*_0_ and at fixed activated-enzyme concentration *E*_0_. We compare these progress curves to the Michaelis-Menten model. A best-fit routine is applied to extract the catalytic turnover rate *k*_cat_ and the Michaelis constant *K*_M_. As an example, **Figure 4A** shows a set of progression curves we acquired using LbCas12a complexed with a guide RNA targeting the wild-type 20 bp ssDNA sequence of the SARS-CoV-2 E gene.^28^ We fixed the activated-enzyme concentration at *E*_0_ = 2 nM and varied the initial uncleaved reporter concentration over values of *S*_0_ from 13 to 800 nM using two-fold serial dilutions. We dispensed each condition in triplicate, sealed the plate, and acquired fluorescence signals at 37 °C in 1× reaction buffer.

**Figure 4.**
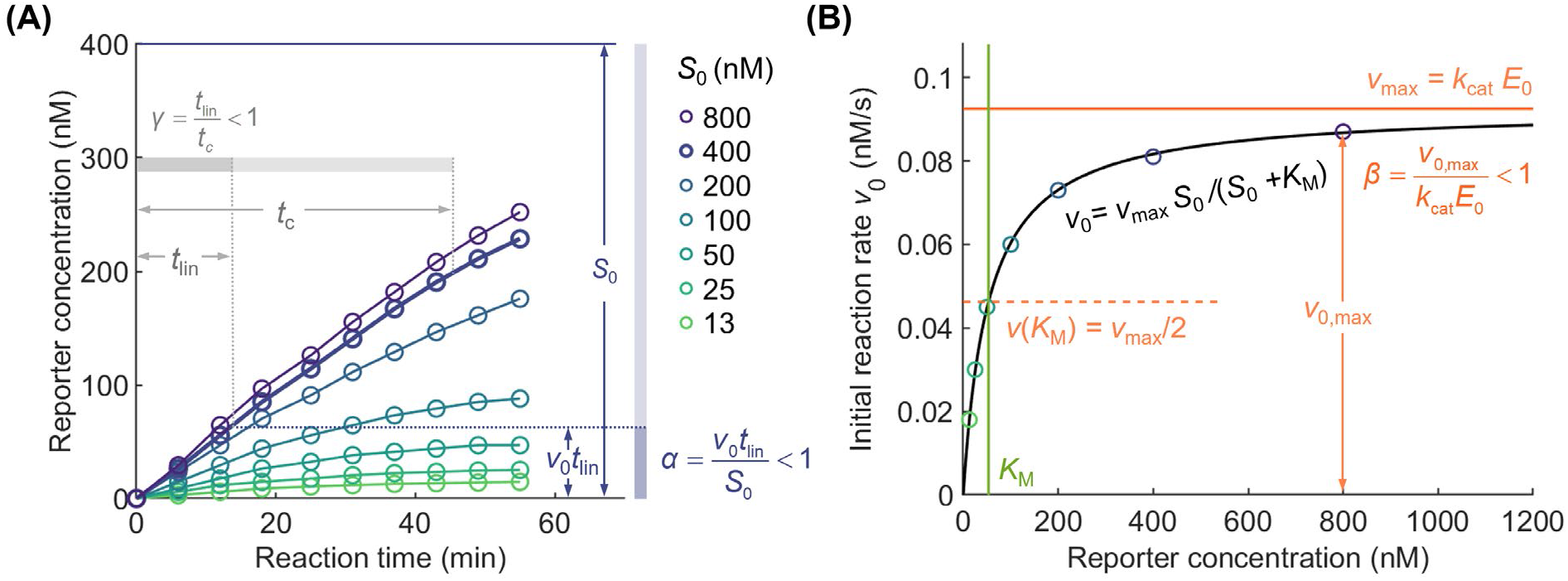
Extraction of Michaelis-Menten kinetic parameters from progress-curve data, with graphical interpretation of the back-of-the-envelope checks *α, β*, and *γ*. **(A)** Cleaved-reporter concentration as a function of time for seven values of the initial substrate concentration *S*_0_ ranging from 13 to 800 nM, at fixed activated-enzyme concentration *E*_0_ = 2 nM. The slope of each curve at early times defines the initial reaction velocity *v*_0_ (*S*_0_).^28^ **(B)** Initial reaction velocities *v*_0_ from (A) plotted against *S*_0_ (symbols) and the Michaelis-Menten fit (solid line, Eq. 3.4) using nonlinear least squares. The fit yields *k*_cat_ = 0.046 s^−1^ and *K*_M_ = 53.3 nM. *α* (blue), *β* (orange), and *γ* (grey) are visualized as annotated colored bars in (A) and (B), where the ratio of the darker bar (numerator) to the lighter bar (denominator) gives the value of each dimensionless number. *t*_lin_, *v*_0_, *v*_0,*max*_ is read directly off the linear early-time portion of the progress curves in (A), while *t*_*c*_ and *v*_max_ = *k*_cat_ *E*_0_ is computed from the best-fit kinetic parameters.

Generally, an experimentalist should strive to obtain progress curves for *S*_0_ values which are both significantlly smaller and significantly higher than *K*_M_. Hence, the range *S*_0_ should be selected based on an educated guess of *K*_M_, or the entire process may be partly iterative. Several general features of progress curves acquired at varying *S*_0_ are important to note. First, the conservation of reporter molecules obviously requires that *P*(*t*) approach but never exceed *S*_0_. Second, progress curves at the low *S*_0_ values can tend to overlap with one another because the rise in signal can become dominated by reporter degradation and detector noise. For example, note the similarity of the experimental progress curves for *S*_0_ =13 and 25 nM in **Figure 4A**. Third, progress curves at *S*_0_ values sufficiently higher than *K*_M_ become approximately independent of *S*_0_ and so exhibit similar initial slopes. For example, the curves at *S*_0_ = 400 and 800 nM in **Figure 4A** exhibit similar slopes at early times.

We recommend progress curves such as **Figure 4A** be analyzed in two steps. First, for each progress curve, we extract the initial reaction velocity *v*_0_ using a simple linear regression fit to the early, linear portion of the curve. These *v*_0_ values versus *S*_0_ are then used to generate the Michaelis-Menten curve. For most Michaelis-Menten analyses, this is then compared to the following simplified initial reaction rates predicted by Michaelis-Menten theory,

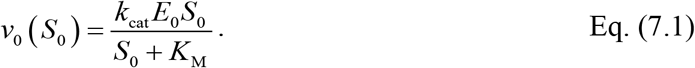

See Section 3 above as well as Avaro and Santiago^13^ for a detailed description of the assumptions behind this equation as well as the regimes of its applicability. In comparing the Michaelis-Menten curve to this theory, we recommend using a nonlinear least-squares regression^24^ to obtain the best-fit values of *v*_max_ and *K*_M_. We provide in **Section S5** of the SI a source code for such a fit. As an example, we show in **Figure 4B** measured *v*_0_ plotted against *S*_0_ (symbols) together with the Michaelis-Menten model fit (solid line). The extracted best-fit parameters are *k* _cat_ = *v* _max_ / *E*_0_ = 0.046 s^−1^ and *K*_M_ = 53.3 nM. The annotated horizontal line at *v* _max_ gives the asymptote of the model. The dashed line at *v* (*K*_M_) = *v*_max_ / 2 provides a graphical reference for *K*_M_, since the abscissa of the half-maximum velocity is *K*_M_. Again, at this point, we recommend confirming that the *S*_0_ values chosen span both well below and well above the measured *K*_M_. As described by Avaro and Santiago^20^, this ensures that the experimental results are sufficiently sensitive to both *k*_cat_ / *K*_*m*_ and *K*_M_. At the same time, the highest chosen values of *S*_0_ should be chosen to avoid the regime of the calibration curve that becomes problematic due to IFE. As described in Section 6.1, for high *S*_0_, the calibration curve strongly deviates from linear and therefore signals become insensitive to *S*_0_ (and so useless). For even higher *S*_0_, the calibration curve eventually double valued (c.f. **Figure 3A**). In **Figure 3A**, this problematic region begins for *S*_0_ above about *c* = 0.3 *C*_0_.

## 8 Check for consistency in own data using α, β, γ: Back-of-the-envelope sanity checks

As first described by Ramachandran and Santiago^18^ and discussed in further detail by Avaro and Santiago^13^, the experimental data in the studies that report the largest kinetic efficiency (i.e., highest values of *k*_cat_ / *K*_M_) often cannot be reconciled with the reported values of the kinetic rates. These inconsistencies point to gross errors in the determination of the kinetic rates of the Cas12 and Cas13 enzymes.^18^ Based on the three published corrections and errata, we attribute most of these errors in the calibration of the fluorescence signal or in unit conversion. Proper calibration is particularly critical for the estimation of *k*_cat_ since this requires an estimation of the maximal reaction velocity in units of concentration per unit time. We therefore recommend that all reports of Michaelis-Menten kinetic rates perform simple, back-of-the-envelope checks for self-consistency between experimental measurements on one hand and the reported kinetic rate parameters *k*_cat_ and *K*_M_ on the other.

Ramachandran and Santiago^18^ first introduced three simple-to-compute ratios for self-consistency which they termed *α, β*, and *γ*. Avaro and Santiago^13^ later proposed a more generally applicable version of *γ* which we will adopt here (and which is valid for a wider range of initial substrate concentrations). The three parameters offer a method to check for self-consistency of the data (and calibration) and to ensure that the reported kinetic rates are consistent with this data. **Figure 4** presents a graphical interpretation of *α, β*, and *γ* upon which we will expand here. The experimental data shown in **Figure 4** are from Blanluet at al.,^28^ who reported Michaelis-Menten *trans*-cleavage kinetic measurements for LbCas12a activated by ssDNA targets.

The check for self-consistency of the enzyme kinetics analysis is based on a check that *α, β*, and *γ* are all less than unity, as dictated by conservation of species and the Michaelis-Menten framework. The first ratio, *α*, follows simply from conservation of reporter molecules. The cleaved-reporter concentration accumulated during the initial, linear regime of a progress curve cannot exceed the initial uncleaved reporter concentration *S*_0_. Hence,

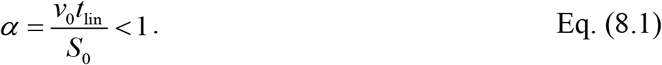

Here *v*_0_ is the measured initial reaction velocity (e.g., in nM s^−1^) and *t*_lin_ is the duration of the initial linear regime. The numerator *v*_0_*t*_lin_ estimates the cleaved-reporter concentration accumulated during the linear regime, and *S*_0_ is the highest cleaved reporter concentration that was used in the Michaelis-Menten study. Here, *t*_lin_ can be estimated either by a visual inspection of the progress curve or, more reproducibly, by fitting a straight line to the first *N* points of the curve. One approach is to use the largest *N* for which the coefficient of determination *R*^2^ remains above a threshold, such as 0.95. This then defines *t*_lin_. The slope of the corresponding linear regression fit is then the estimate of *v*_0_. Importantly, the estimation of *α* doesn’t require knowledge of the kinetic rate parameters. **Figure 4A** illustrates a graphical interpretation where *α* is depicted as fraction of the total *S*_0_ magnitude (denoted by a vertical bar). In the illustration, dark blue band has length *v*_0_*t*_lin_ while the full blue band has length *S*_0_.

The second ratio, *β*, compares the measured initial reaction velocity to the theoretical maximum velocity predicted by the Michaelis-Menten model,

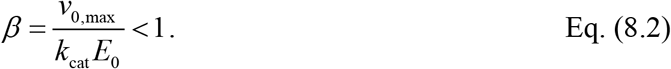

Here *v*_0,max_ is the maximum measured initial reaction velocity across the *S*_0_ dilution series. *v*_0,max_ corresponds to the highest *S*_0_ value used in the experiments. This can be determined from the corresponding progress curve (c.f. **Figure 4A**) or from the high *S*_0_ asymptote of the Michaelis-Menten curve (c.f. **Figure 4B**). Of course, the asymptotic reaction velocity *v*_max_ = *k*_cat_ *E*_0_ is the theoretical maximum reached in the limit *S*_0_ ≫ *K*_*M*_. Comparing the experimentally measured maximum reaction velocity from the progress curves or Michaelis-Menten fit with the theoretical value provides a consistency check for the reported values of *k*_cat_ and *E*_0_. *β* is graphically depicted in **Figure 4B**.

The third ratio, *γ*, compares an estimate of the linear time scale *t*_lin_ to the characteristic completion time of the *trans*-cleavage reaction *t*_c_. We can define *t*_c_ as the time at which *S* (*t*_c_) / *S*_0_ = *e*^−1^. Starting from a closed-form solution of the Michaelis-Menten model, Avaro and Santiago^13^ derived and introduced the following generalized definition of *t*_c_:

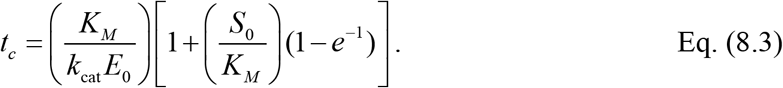

Therefore, we can define our check as

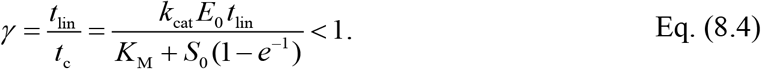

In the low-*S*_0_ limit (*S*_0_ ≪ *K*_*M*_), *t*_c_ reduces to *K*_M_ / (*k*_cat_ *E*_0_) (the value suggested by Ramachandran and Santiago^18^). In the high-*S* limit (*S*_0_ ≫ *K*_*M*_), *t*_c_ reduces to *S* (1− *e*^−1^) / (*k* _kat_ *E*_0_). The limit for *γ* < 1 described by Eq. 8.4 is valid for both regimes. **Figure 4A** shows a graphical interpretation of *γ* as the ratio of two timescales. Unlike *α, γ* cannot be read directly off the progress curves alone. *t*_c_ is computed entirely from the reported values of *k*_cat_ and *K*_M_, and known values of *E*_0_ and *S*_0_. The estimate for *t*_lin_ was described above.

A practical note on dimensions of relations and units of reported quantities is that *α, β*, and *γ* are non-dimensional only if all concentrations and all rate constants are expressed in consistent units. We recommend converting all concentrations to nM and all rate constants to s^−1^ before computing the three ratios. From our reviews of papers which have inconsistencies,^13^ a common pitfall is using incorrectly comparing concentrations and mixing seconds and minutes. For example, in one study^42,43^, errors in each of these led to an over reporting of *k* by 6 orders of magnitude. Two of the current authors communicated this to these researchers and they updated their *k*_cat_ from 3.9 × 10^5^ s^−1^to 0.24 s^−1.13,42,43^

We provided a **Script 3** (c.f. Section S4 in SI)^41^ that is useful in performing the complete pipeline of Sections 7 and 8. This includes calibration, to progress curves, to extraction of *k*_cat_ and *K*_M_, and to computation of *α, β*, and *γ*. Commercial software packages such as GraphPad Prism (GraphPad Software, San Diego, CA) and OriginPro (OriginLab, Northampton, MA) provide alternatives for fitting the Michaelis-Menten model.

### Step-by-step protocol

#### Acquire progress curves at varying initial substrate concentrations

1. Prepare a serial dilution of intact uncleaved reporter in 1× reaction buffer over seven concentrations spanning two orders of magnitude. Keep the highest concentration below the approximate IFE boundary where the calibration is useful: *c*\* < 0.3 (Section 6.1).
2. Activate the CRISPR ribonucleoprotein by incubating the gRNA-Cas complex with the target at 37 °C for 30 min.
3. Mix activated RNP with each reporter dilution to a final activated-enzyme concentration *E*_0_ identical across all wells and dispense each condition into triplicate. Use the same well volume as in Sections 5 and 6.
4. Seal the plate, briefly centrifuge to remove bubbles, and acquire fluorescence using the same instrument settings and sampling interval as the calibration acquisitions. Acquire for a duration sufficient to capture saturation of the progress curves, typically 60 min.
5. Apply the flat-field and background correction (Eq. 5.1) and the IFE-corrected calibration (Eq. 6.4) to convert the raw fluorescence to *P*(*t*) at each well and time point.

#### Extract initial reaction velocities and fit the Michaelis-Menten model

1. For each *S*_0_, fit a linear regression line to the initial, linear portion of the curve. One method of determining this is to fit the first *N* data points of a few curves and then increase *N* until *R*^2^ first drops below 0.95. We then choose a reasonable single value of *t*_lin_ which we apply to all curves. The slopes of the linear fits are *v*_0_.
2. Plot *v*_0_ against *S*_0_ and fit the data to Eq. (7.1) to obtain *v*_max_ and *K*_M_. Compute *k*_cat_ = *v*_max_ / *E*_0_ and report 95% confidence intervals.

#### Compute *α, β, γ* and report

1. Convert all concentrations to a single molar unit (e.g., nM) and all rate constants to s^−1^ before computing the three ratios.
2. Compute *α, β*, and *γ* from Eqs. (8.1, 8.2, 8.4) for each *S*_0_, and verify that all three are less than unity.
3. We provide an open-source implementation^41^: see **Script 3** in SI, which takes *P*(*t*), *E*_0_, and the array of *S*_0_ as input and returns *k*_cat_, *K*_M_, *t*_lin_, *v*_0_, *α, β*, and *γ*.

## 9 Determination of LoD

Sections 7 and 8 focused on the quantification of kinetic rate parameters in the Michaelis-Menten framework. We here focus on experiments useful in the design of CRISPR-based assays. Specifically, the measurement of assay LoDs. We will define LoD simply as the lowest target concentration that produces a signal distinguishable from a no-target control (NTC). It defines the clinical or analytical range over which the assay is useful. Also, the LoD is the most widely reported figure of merit for diagnostic assays and is used to compare CRISPR-based assays against alternative methods such as PCR and LAMP.

Reviews of the CRISPR diagnostics literature reported that LoDs are often poorly characterized.^11,14,44,45^ In particular, CRISPR assays without pre-amplification (e.g., using PCR or LAMP) can be very difficult to reconcile with the low kinetic rates of CRISPR enzymes. Hence, assay design should involve both quantification of kinetic rates (Sections 6 and 7) and some determination of LoD, which we present here.

As described by Huyke et al.,^28^ two complementary experimental readouts of the LoD have been used in the literature: an endpoint-based LoD, *L*_e_, and a rate-based LoD, *L*_v_. *L*_e_ is defined as the smallest target concentration *E*_0_ for which the cleaved-reporter concentration *P* (*t*) at a chosen endpoint time *t* exceeds a threshold determined by the NTC (and its repeatability). *L*_v_ is defined as the smallest *E*_0_ for which the initial reaction velocity *v*_0_ exceeds a threshold set by the NTC velocity threshold (and its repeatability). Following Huyke et al.,^17^ we will here set these thresholds as three standard deviations above the NTC mean.

The LoD of any CRISPR assay is fundamentally limited by the background signals of the no-target control (NTC). In the absence of target, Avaro and Santiago^22^ showed that, for fluorescence-based CRISPR assays, the background signal is most often dominated by temperature-dependent degradation of reporter molecules in the solution. Specifically, using a simple first-order kinetics framework for this degradation, the ability to resolve initial CRISPR *trans-*cleavage reaction velocities depends solely on reporter degradation, *v*_BG_ = *k*_rep_ *S*_0_ (c.f. Section 6.2). The kinetic rate can be very low, but this low rate can dominate at trace target concentrations. For example, Avaro and Santiago^22^ showed reporter chemistry used exhibits an *k*_rep_ of order 10^−6^ s^−1^, but this low rate can dominate signal for target concentrations below about 1 pM. Hence, we can define a background velocity set by the reporter chemistry alone. Importantly, this value is then independent of the CRISPR enzyme or target. An assay can detect target only when the *trans*-cleavage signal driven by activated enzyme exceeds this degradation background.

Avaro and Santiago^22^ developed a theoretical framework to obtain a closed-form expression for the rate-based LoD. This framework leads to the following estimates for *L*_v_,

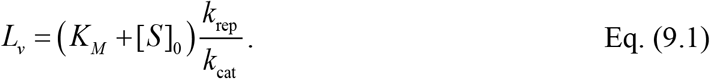

This equation shows that the minimum detectable target is governed by a ratio of (undesired) reporter cleavage and the (desired) enzyme-specific *trans-*cleavage.

We next discuss an estimate for *L*_e_. To this end, we here extend the analysis of Avaro and Santiago^22^ to derive a closed-form expression for this endpoint LoD, which is more commonly used. Following the same augmented Michaelis-Menten model^22^, we equate the corrected fluorescence signal in the presence of target with the corrected signal in the no-target control at a chosen endpoint time *t* (see Section S3 in SI for the full derivation). This yields the theoretical endpoint LoD,

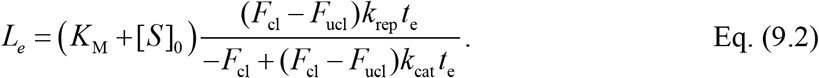

Where *F*_Cl_ and *F*_Ucl_ are the calibration parameters of Section 6.1. This relation also highlights the importance of a judicious choice of endpoint time for an assay. Specifically, *L*_e_ decreases monotonically with the endpoint time *t*_e_, and this captures the tradeoff between assay duration and assay sensitivity. For endpoint times *t* ≪ *F*_Cl_ / [*k*_cat_ ×(*F*_Cl_ − *F*_Ucl_)], *L*_e_ of Eq. (9.2) approaches the asymptote *L*_v_ of Eq. (9.1).

We note that the theoretical rate-based LoD is the lower limit of the endpoint LoD, and we have *L*_v_ < *L*_e_ for typical assay times between 15 to 60 min. The rate-based readout is therefore here recommended to achieve a lower (better) LoD, especially for shorter assays. Rate-based readout has the additional advantage that it does not require an assay designer to select an a priori a specific time (or times) to evaluate the assay results.

We stress that Eqs. (9.1) and (9.2) are informative even before any experiment is performed. For these estimates highlight the importance and utility of obtaining an estimate of *k*_rep_ (c.f. Section 6.2) and having at least estimates of enzyme kinetics (Section 7). For example, LoD scales linearly with *k*_rep_ and inversely with the catalytic efficiency *k*_cat_ *K*_M_ in the limit of *S*_0_ ≪ *K*_*M*_. This also highlights areas for future research and improvements in CRISPR assay systems: improving (decreasing) LoD requires either a more stable reporter (lower *k*_rep_) or a more catalytically efficient enzyme (higher *k*_cat_ / *K*_M_), or both. Lastly, note that reporter concentration *S*_0_ also affects the LoD. For example, Eq. (9.1) shows that *L*_v_ scales linearly with *S*_0_ for *S*_0_ ≫ *K*_*M*_. In this regime, increasing reporter concentration might increase catalytic rates but at the great expense of adding background and decreasing the LoD. Recall also that in Section 6.1 we pointed out that lowering *S*_0_ is also important in resolving the linear regime of the calibration curve. An accurate calibration curve can itself improve LoD.

**Figure 5** shows an example LoD determination using LbCas12a based on the experimental data of Huyke et al.^28^ using ssDNA activator. The top row labeled “DATA” shows measurements of cleaved-reporter concentration versus time (progress curves) at seven activator concentrations from 10 fM to 10 nM in factor-of-ten steps at fixed *S*_0_ = 200 nM, in addition to NTC experiments. The plots in the middle row labeled “READOUT” show the rate-based readout *v*_0_ (*E*_0_) (left) and the endpoint readout *P* (*E*_0_;*t* = *t*_e_) (right). The endpoint time for the latter was *t*_e_ *=* 50 min. Three operating regimes are observed. At activator concentrations of 1 pM and below, the progress curves overlap with the NTC. Here, the *trans*-cleavage signal is below the reporter degradation background so signal is dominated by reporter degradation. At 10 to 100 pM, the progress curves separate from the NTC and define the usable dynamic range. At 1 nM and above, the progress curves saturate within the assay window, and both the endpoint signal and the initial reaction velocity under-report true target abundance. We refer to the latter regime as the late-readout regime. Avaro and Santiago^22^ presented a closed-form estimate of the onset of this regime as *E*_0_ > *K*_M_ / (*k*_cat_*τ* _pre_), where *τ* _pre_ is a time scale associated with mixing of the reagents (e.g., pipetting into the microwell plate) and loading and startup of the thermocycler instrument. In each panel, the dashed horizontal line labeled “Degradation floor” marks the NTC threshold. The solid line curve is a four-parameter logistic fit ranging from 0.1 pM to 1 nM. The intersection of the two dashed lines and shaded rectangle marks the intersection between the logistic fit and the threshold. This intersection point is labeled with a star symbol and defines the experimental values of LoD *L*_e_ and *L*_v_. The bottom row labeled “THEORY” shows the closed-form predictions for *L*_e_ from Eq. (9.2) and *L*_v_ from Eq. (9.1). The experimentally determined *L*_v_ is lower than *L*_e_ on the same dataset, consistent with the inequality predicted by the theoretical framework.

**Figure 5.**
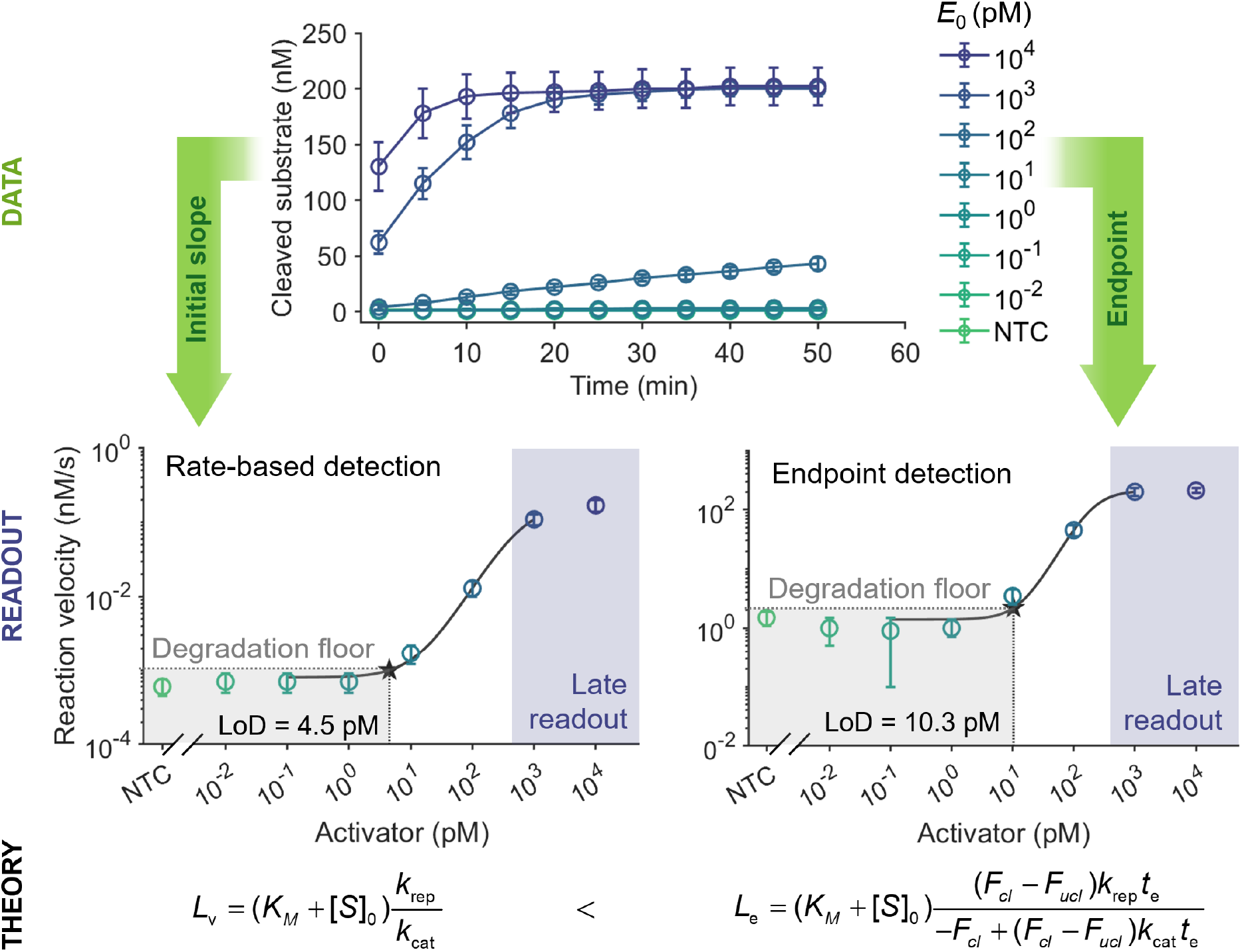
Determination of the limit of detection (LoD) of a CRISPR assay by titration of the activated enzyme concentration *E*_0_. The top plot shows example measured cleaved-substrate progress curves for values of *E*_0_ ranging from 10 fM to 10 nM, and a no-target control (NTC), all at fixed initial substrate concentration of *S*_0_ = 200 nM. The progress curves of lowest *E*_0_ overlap with the NTC, illustrating the regime where reporter degradation limits signal. The highest *E*_0_ traces saturate within the assay duration, illustrating the so-called late-readout regime where measured endpoint values and reaction velocities under-report the true initial conditions of the trans-cleavage reaction. The two lower plots show two determinations of the LoD based on signal increase rate (left) and endpoint readout (right). On the left is plotted initial reaction velocity as a function of activator concentration. On the right is the final signal after fixed reaction times (here at *t*_e_ *=* 50 min) as a function of activator concentration. In each panel, the dashed horizontal line marks the reporter degradation floor limited by *k*rep. A four-parameter logistic fit (solid line) intersects degradation floor at the LoD (star). The shaded regions annotate the degradation-limited regime (grey, low *E*_0_) and the late-readout regime (purple, high *E*_0_). Along the bottom are theoretical, closed-form expressions for the two LoDs. One for rate-based LoD, *L*_v_, and a second for endpoint LoD, *L*_e_. The inequality between the two analytical expressions confirms that theoretical rate-based LoD is always lower than the that of the endpoint LoD, consistent with the experimental results shown in the middle row.

Lastly, we offer one practical note on titration of target. For low target concentrations (e.g., sub-picomolar concentrations), adsorption of the target nucleic acid to the wetted walls of the well plastic can deplete the working concentration (activity) relative to the dispensed concentration. For such low concentrations, we recommend pre-coating the wells with carrier nucleic acid or a non-specific blocker (e.g., poly-A oligonucleotides, BSA) when working below 10 pM target.^46^ We also recommend that target serial dilutions be performed in low-binding tubes and used immediately after preparation.

### Step-by-step protocol

#### Acquire progress curves at varying activator concentrations

1. Prepare a serial dilution of the target nucleic acid (activator) in 1× reaction buffer over at least seven concentrations spanning 4 to 6 orders of magnitude and a no-target control (NTC) prepared in the same buffer without activator.
2. For activator concentrations below 10 pM, pre-coat the wells with a non-specific blocker (e.g., 0.1 % BSA or carrier poly-A oligonucleotide) to reduce surface adsorption losses of the target.^46^ Prepare serial dilutions in low-binding tubes and use them immediately after preparation.
3. Activate the CRISPR ribonucleoprotein by incubating the gRNA-Cas complex with the target at 37 °C for 30 min.
4. Mix the activated RNP with the uncleaved reporter at fixed *S*_0_ identical across all wells. Keep the reporter concentration under the IFE suggested boundary of *c* < 0.3 *C*_0_ (Section 6.1).
5. Dispense each activator condition into triplicate wells of the same plate. Use the same well volume as in Sections 5 through 8.
6. Seal the plate, briefly centrifuge to remove bubbles, and acquire fluorescence at 37 °C using the same instrument settings and sampling interval as the calibration acquisitions of Sections 5 and 6. Acquire for the chosen assay duration, typically 15 to 60 min.

#### Process raw fluorescence to obtain *P*(*t*)

1. Apply the flat-field and background correction (Eq. 5.1) and the IFE-corrected calibration (Eq. 6.4) to convert the raw fluorescence to the cleaved-reporter concentration *P*(*t*).
2. Average the triplicate progress curves for each activator condition. Compute the standard deviation across the NTC triplicates for use in the threshold calculations below.

#### Compute the rate-based LoD *L*_v_

1. For each activator condition and the NTC, extract the initial reaction rate from the averaged progress curve using linear regression (following the same process in Section 8).
2. Compute the NTC velocity threshold as the mean of the NTC *v*_0_ values plus 3 times the standard deviations.
3. Plot *v*_0_ as a function of *E*_0_ on a log-linear axis. Fit a four-parameter logistic curve to the data.
4. *L*_v_ is the *E*_0_ at which the logistic fit crosses the NTC velocity threshold.

#### Compute the endpoint-based LoD *L*_e_

1. Choose an endpoint time *t*_0_ within the assay window. We typically use *t*_0_ between 15 and 60 min.
2. Record and report *P*(*t*_0_) for each activator condition.
3. Compute the NTC endpoint threshold as the mean of the NTC *P*(*t*_0_) values plus 3 times the standard deviations.
4. Plot *P*(*t*_0_) as a function of *E*_0_ on a log-linear axis. Fit a four-parameter logistic curve to the data.
5. *L*_e_ is the *E*_0_ at which the logistic fit crosses the NTC endpoint threshold.

#### Compare with theoretical LoDs

1. Compute the theoretical rate-based LoD, *L*_v_, from Eq. (9.1) using *k*_cat_ and *K*_M_ from Section 7, *k*_rep_ from Section 6.2, and the chosen *S*_0_.
2. Compute the theoretical endpoint-based LoD, *L*_e,_ from Eq. (9.2) at the chosen *t*_0_, using the calibration parameters *F*_Cl_ and *F*_Ucl_ from Section 6.1.
3. Compare the experimentally determined *L*_v_ and *L*_e_ with their theoretical predictions. A large discrepancy may indicate an error in the calibration (Sections 5 and 6) or the kinetic-rate measurement (Section 7).
4. We provide an open-source implementation^41^: see **Script 4** in SI, which takes the calibrated progress curves *P*(*t*) at varying activator concentrations, *k*_cat_, *K*_M_, *k*_rep_, *F*_Cl_, and *F*_Ucl_ as input and returns *L*_v_, *L*_e_, and their theoretical predictions from Eq. (9.1) and Eq. (9.2).

## 10. Comments regarding enzyme assays performed with miniaturized compartmental reaction systems

The current paper has focused on bulk fluorescence measurements acquired using a thermocycler or plate reader (c.f. Section 4). Such readout systems constitute the vast majority of studies reporting enzyme kinetics of CRISPR systems, and such systems are very commonly used to demonstrate and develop new CRISPR-based assays. However, some of the concepts we discuss here may be applicable to other reaction formats. We here offer comments on CRISPR kinetics studies and CRISPR diagnostic assay development using microfluidic systems where many parallel reactions are performed.^11,47^ The two most widely adopted formats are microfluidic droplet systems ^48,49^ and specialized micron-scale microwell chamber devices.^50,51^

Droplet digital CRISPR (ddCRISPR) formats partition a bulk sample and the necessary CRISPR reagents into tens of thousands of independent, water-in-oil microdroplets, typically in 𝒪(10-100 pL) volume range. This physical partitioning can isolate individual target molecules within droplets and can be used to reduce competition among target-containing compartments. For example, Shan et al.^48^ used droplet microfluidics to isolate single SARS-CoV-2 RNA recognition events. Such platforms have been used to study amplification-free LoDs in the sub-femtomolar range.

Similarly, femtoliter chamber-array platforms compartmentalize CRISPR reactions into millions of microreactors fabricated using microfabrication methods such as fluoropolymer photolithography or injection-molded plastics such as COP and polycarbonate.^51^ Amplification-free CRISPR detection using femtoliter chamber arrays are very likely the highest sensitivity implementations of CRISPR assays with limits of detection as low as 6.5 aM for viral RNA.^52^ In the latter work, CRISPR Cas13 assay reagents and sample were dispensed using specialized methods into arrays of microwells and fluorescence signals are obtained by rapid optical scanning of compartments.^50^ Incorporation of a target-capture step using magnetic bead-based enrichment was used to improve assay sensitivity 7-fold.

As pointed out by Avaro and Santiago^13,14^, the extremely high sensitivity of micron-scale compartmentalized formats may be due to the molecular confinement. Consider that a single activated enzyme molecule confined within a 1 fL microwell is equivalent to a local enzyme concentration of approximately 1.7 nM. The latter is an effective concentration much higher than the 𝒪(1 pM) of the original solution. The confinement also restricts cleaved reporters to a small volume, increasing signal intensity.

While compartmentalization can significantly improve assay sensitivity, it also introduces additional optical and analytical considerations for quantitative fluorescence measurements. First, consider the flat field correction discussed in Section 5. We hypothesize this correction may be important for such micro-compartmentalized systems. For example, consider the system of Iida et al.,^50^ which obtains fluorescence readouts from up to 10^5^ to 10^6^ femtoliter chambers within a single field of view. They addressed signal nonuniformity through per-chamber background subtraction and fluorescence calibration. A second example is the work of Shan et al.^48^, which utilizes wide-field fluorescence imaging to read out thousands of CRISPR-containing microdroplets. These systems may require correction for non-uniform illumination due to vignetting, stray light, and position-dependent lens transmission.

Second, the current discussions of calibration (Section 6) should apply to any systems where optical signals are used to deduce product species concentrations in enzymatic reactions. For example, the workflow of Shinoda et al.^51^ first calibrates in-chamber fluorescence against known reporter concentrations before extracting quantitative *trans*-cleavage data. The calibration guidelines provided here should be applicable. One important distinction of very small volumes is that their small associated optical pathlengths (ranging from ~10 to 100 µm for droplets, to ~1 to 5 µm for microwell chambers) are unlikely to require correction for IFE.

Third, miniaturization of droplets and femtoliter chambers greatly increases the surface-to-volume ratio compared to standard well plates. Hence, surface effects may be exacerbated. For example, Cas enzymes, crRNA, and/or reporter molecules may adsorb to solid chamber walls or to the water-oil interface of droplets, reducing the activity of reagents and therefore biasing measurement. The effect is generally stronger in smaller compartments such as femtoliter chambers than in larger picoliter droplets. To reduce surface interactions, reaction buffers have included passivation agents and/or surfactants such as bovine serum albumin (BSA) and Triton X-100. For example, Shinoda et al.^51^ included Triton X-100 in the reaction buffer of a femtoliter chamber-array CRISPR system to suppress non-specific surface interactions.

## 11. Summary

Quantitative fluorescence-based enzyme assays require rigorous experimental methods for signal calibration, kinetic analyses, and interpretation of signal relative to background noise. In CRISPR-Cas diagnostic assays, these requirements are particularly important because fluorescence calibration directly affects the reported enzyme kinetic rates and assay limits of detection. The motivation of this work was therefore to provide a practical and quantitative framework for reproducible fluorescence-based CRISPR enzyme assays using standard laboratory instrumentation.

We introduced a fairly comprehensive framework and step-by-step protocols for the quantification, evaluation, and development of CRISPR enzyme assays that use fluorescence readout. We first reviewed the relevant Michaelis-Menten framework and its relevance to reaction rates. We then presented a detailed process for signal calibration. This included flat-field and background correction of the raw fluorescence images, and IFE-corrected calibration of both cleaved and uncleaved reporters. After calibration, we considered two major workflows. The first was to quantify enzyme reaction kinetics. This is critical because the *trans-*cleavage reaction rates of CRISPR-Cas enzymes govern the sensitivity of assays that do not rely on pre-amplification with techniques such as PCR and LAMP. The second workflow was focused on quantitative methods to develop and evaluate assays with high sensitivity and to measure an assay’s limit of detection.

For the first workflow, we described experimental protocols to generate progress curves and ultimately generate a Michaelis-Menten curve. We provided guidelines for obtaining Michaelis-Menten curves that are sensitive to key parameters, and for fitting experimental data to extract enzyme kinetic parameters. We summarized simple, back-of-the-envelope relations that can be used to check for self-consistency in CRISPR kinetics data. For the second workflow, we provided methods for quantitative determination of background signals and reaction rates, with an aim at optimizing limits-of-detection of CRISPR-Cas nucleic acid assays. As part of this, we contrasted and quantified detection based on either initial rate or endpoint fluorescence.

Lastly, we stressed that rational and repeatable experimental methods are essential to advancing CRISPR-Cas enzyme diagnostic assays and their implementation. In particular, explicit and correct signal calibration is crucial for the accurate determination of both enzyme kinetics and assay limit of detection. Reporting these calibration steps alongside the kinetic rates and LoD measurements is essential for comparability across studies and for enabling other labs to reproduce results. We hypothesized that errors in this calibration step have largely led to variations of more than 6 orders of magnitude for the reported value of *k* _cat_ across labs for the same Cas ortholog.^13^

To facilitate the adoption of the methods presented in this work, we provided open-source Python implementations for every step in this protocol and encourage the community to adopt shared tools and standard reporting templates of this kind. Such standardization and refinement of experimental protocols should support the continued development and broader translation of CRISPR diagnostic technologies.

