## Supporting Information file for "Experimental Methods for CRISPR Enzyme Assays with Fluorescence Readout"

### Contents

S1 Recommended Reagents and Equipment

S2 Evaluation of flat-field and background correction

S3 Derivation of theoretical limit of detection (LOD)

S4 Open-source Jupyter Notebook script

**S1 Recommended Reagents and Equipment**

We provide additional details on the reagents, instruments, and experimental considerations typically used for and relevant to fluorescence-based CRISPR enzyme assays. The section summarizes recommended CRISPR enzymes, oligonucleotides, buffers, fluorescence reporters, thermocyclers, and fluorescence detection systems compatible with quantitative fluorescence measurements and calibration. These materials and platforms provide a practical starting point for establishing fluorescence-based CRISPR assays and support reproducible kinetic measurements and fluorescence signal analysis.

**Reagents:**

- EnGen® Lba Cas12a (Cpf1) (New England Biolabs, cat. no. M0653T). LbCas12a (cat. no. 10007922) and AsCas12a (cat. no. 10001272) can also be purchased from Integrated DNA Technologies, or prepared using published plasmid sequences.
- NEBuffer r2.1 for Cas12a assays (cat. no. B6002S). Optimal buffers vary with Cas ortholog.
- Oligonucleotides, including guide RNA (gRNA), synthetic target DNA, and fluorophore-quencher probes can be obtained from different sources (e.g., Integrated DNA Technologies, Elim Biopharmaceuticals Inc., GeneLink Inc., or Eurofins Genomics).
- Nuclease-free water

**Equipment:**

- Air displacement pipette
- Pipette tips
- Thermocycler and/or fluorescence microtiter plate reader. The following protocol has been tested using MiniOpticon thermal cycler (Bio-Rad Laboratories, CA, USA) and a CFX384 Touch Real-Time PCR Detection System (Bio-Rad Laboratories).
- Hard-Shell® 384-Well PCR Plates, thin wall, skirted, clear/white (cat. no. HSP3805, Bio-Rad Laboratories) to use with the CFX384 system; other well plates or tube strips for fluorescence readout which can contain 30 to 50 µl and are compatible with the thermocycler we used.

### S2 Evaluation of flat-field and background correction

In this section, we quantify the effect of flat-field and background correction on well-to-well fluorescence variability and evaluate the temporal stability of the correction during kinetic measurements.

**Figure S1(A)** shows the coefficient of variation (CV) across triplicate wells for seven reporter concentrations spanning 14 to 900 nM, before (black circles) and after (orange squares) flat-field and background correction. The bars show the corresponding mean corrected fluorescence signal at each concentration on a logarithmic scale. The correction decreases the within-triplicate CV for six of seven concentrations, with the largest reductions (~1%) observed at intermediate reporter concentrations of 225 and 450 nM. This reduction is consistent with removal of position-dependent systematic bias arising from spatial nonuniformity in illumination, optics, and detector response. We hypothesize that the residual CV primarily reflects random pipetting error and detector noise. At the highest reporter concentration (900 nM), the correction slightly increases the CV. In this regime, the raw signal is likely dominated by random variability rather than spatial systematic bias, and the correction introduces a small additional uncertainty associated with the measured flat-field and background matrices.

**Figure S1(B)** shows the temporal evolution of the mean well-to-well CV across reporter concentrations for the raw and corrected fluorescence signals during a representative 2150 s experiment. After the first 250 s, the correction consistently reduces the mean CV from 4.0 % to approximately 3.5 %. The well-to-well CV of the corrected fluorescence signal varies in time. The figure also plots the difference, $\Delta\mathrm{CV} = \mathrm{CV}_{\mathrm{raw}} - \mathrm{CV}_{\mathrm{corrected}}$. At long times ($t > 1500$ s), $\Delta\mathrm{CV} = 0.5\,\%$. This corresponds to an average reduction (improvement) of 0.5% relative to the raw fluorescence signal. We stress that $I_{\mathrm{FF}}(t)$ and $I_{\mathrm{BG}}(t)$ should be acquired on the same temporal grid as $I_{\mathrm{raw}}(t)$.

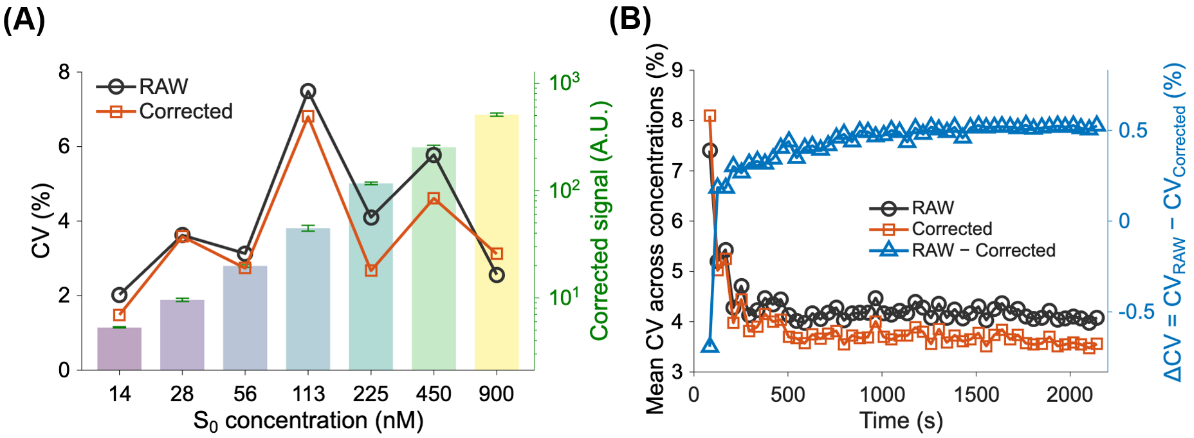


**Figure S1.** Evaluation of flat-field and background correction. (A) Coefficient of variation (CV) among triplicate wells before and after flat-field and background correction. Bars show the corresponding mean corrected fluorescence signal. (B) Temporal evolution of the mean well-to-well CV for raw and corrected fluorescence signals during a representative kinetic experiment. The blue data show the difference $\Delta \mathrm{CV} = \mathrm{CV}_{\mathrm{raw}} - \mathrm{CV}_{\mathrm{corrected}}$.

### S3 Derivation of theoretical limit of detection (LoD)

We here derive the theoretical endpoint limit of detection $L_e$ discussed in Section 9 of the main text. The derivation follows the augmented Michaelis-Menten framework proposed by Avaro and Santiago,[1] together with the fluorescence calibration model introduced in Section 6.1. The theoretical rate-based LoD,

$$L_v = \left(K_M + S_0\right)\frac{k_{\text{rep}}}{k_{\text{cat}}}, \tag{S3.1}$$

was previously derived by Avaro and Santiago.[1] Here, we extend that framework to derive a corresponding closed-form expression for the endpoint LoD.

Starting from the closed-form solution of the Michaelis-Menten progress curve, the corrected fluorescence signal can be written as[1]

$$I_{\text{corr}}(t;E_0) = 10^{-\frac{S_0}{c_0}}\left(F_{\text{ucl}}K_M\mathcal{A}(t) + F_{\text{cl}}\left(S_0 - K_M\mathcal{A}(t) - E_0\frac{\mathcal{A}(t)}{1+\mathcal{A}(t)}\right)\right), \tag{S3.2}$$

where

$$\mathcal{A}(t) = W\left(\frac{S_0}{K_M}\exp\left[\frac{S_0}{K_M} - \frac{k_{\text{cat}}}{K_M}E_0 t\right]\right), \tag{S3.3}$$

and $W$ denotes the Lambert-$W$ function.

We next consider the trace-target limit where

$$\frac{k_{\text{cat}}}{K_M}[E]_0 t \ll 1, \tag{S3.4}$$

Using a first-order expansion, Eq. (S3.2) simplifies to

$$I_{\text{corr}}(t;E_0) \approx 10^{-\frac{S_0}{c_0}} S_0\left(F_{\text{cl}}\left(1 - \frac{E_0}{K_M + S_0}\right) - (F_{\text{cl}} - F_{\text{ucl}})\left(1 - \frac{k_{\text{cat}}E_0 t}{K_M + S_0}\right)\right). \tag{S3.5}$$

To estimate the endpoint LoD, we compare this fluorescence signal with the no-target control. Following the reporter-degradation model of Section 6.2, the corrected fluorescence signal in the absence of target is

$$I_{\text{corr,BG}}(t) = 10^{-\frac{S_0}{c_0}} S_0\left(F_{\text{cl}} - (F_{\text{cl}} - F_{\text{ucl}})e^{-k_{\text{rep}}t}\right). \tag{S3.6}$$

Assuming $k_{\text{rep}}t \ll 1$, we approximate $e^{-k_{\text{rep}}t} \approx 1 - k_{\text{rep}}t$. We define the theoretical endpoint LoD as the target concentration for which the fluorescence increase due to CRISPR *trans*-cleavage equals the fluorescence increase caused by reporter degradation. Equating Eqs. (S3.5) and (S3.6) yields

$$L_e = (K_M + S_0)\frac{(F_{\text{cl}} - F_{\text{ucl}})k_{\text{rep}}t_{\text{e}}}{-F_{\text{cl}} + (F_{\text{cl}} - F_{\text{ucl}})k_{\text{cat}}t_{\text{e}}}. \tag{S3.7}$$

Equation (S3.7) shows that the endpoint LoD decreases monotonically with the endpoint time $t_{\text{e}}$, reflecting the tradeoff between assay duration and assay sensitivity. In the limit

$$t \gg \frac{F_{\mathrm{cl}}}{k_{\mathrm{cat}}(F_{\mathrm{cl}} - F_{\mathrm{ucl}})}. \tag{S3.8}$$

Eq. (S3.7) approaches

$$L_e \to (K_M + S_0)\frac{k_{\mathrm{rep}}}{k_{\mathrm{cat}}} = L_v. \tag{S3.9}$$

The endpoint LoD therefore asymptotically approaches the theoretical rate-based LoD derived by Avaro and Santiago.[1] For experimentally relevant assay times, $L_e$ is always larger (worse) than $L_v$.

### S4 Open-source Jupyter Notebook Script

We provide an open-source Google Colab Jupyter notebook implementing all analysis steps described in this protocol,[2] including flat-field and background correction, fluorescence calibration, Michaelis-Menten kinetic analysis, self-consistency checks, and limit-of-detection (LoD) determination.

The notebook and all associated example datasets are publicly available at:
Github repository: https://github.com/qijiang-tech/cripr-protocol
Google Colab notebook: https://colab.research.google.com/github/qijiang-tech/cripr-protocol/blob/main/crispr_protocol.ipynb

The notebook is organized into one mandatory setup section followed by four largely independent scripts. After running the Setup section, users may execute individual scripts independently depending on the desired analysis workflow. The Setup section imports all required Python libraries and defines helper functions for loading, reshaping, visualizing, and processing fluorescence plate-reader data.
In **Script 1**, flat-field and background correction of raw fluorescence plate-reader measurements is performed using the calibration procedure described in Section 5. The script aligns background, flat-field, and experimental fluorescence arrays, applies the correction to each well at every time point, and exports corrected fluorescence data for subsequent analysis.
In **Script 2**, fluorescence calibration of cleaved and uncleaved reporters is performed using the IFE-aware calibration framework introduced in Section 6. The script extracts the calibration constants for cleaved and uncleaved reporters and determines the characteristic IFE concentration parameter. It also identifies the operational concentration regimes and quantifies the reporter degradation rate constant $k_{\text{rep}}$.
In **Script 3**, progress-curve data acquired at varying substrate concentrations is analyzed to extract Michaelis-Menten kinetic parameters. The script computes initial reaction velocities, fits the Michaelis-Menten model to determine $V_{\text{max}}$ and $K_M$, estimates $k_{\text{cat}}$, and evaluates the self-consistency parameters $\alpha$, $\beta$, and $\gamma$.
In **Script 4**, both endpoint and rate-based limits of detection are analyzed from progress curves measured at varying target concentrations. The script computes experimental LoDs from interpolation of threshold crossings and also evaluates theoretical LoDs using the kinetic and calibration parameters obtained from **Scripts 2 and 3**.

The repository additionally includes example datasets corresponding to each analysis workflow, including representative flat-field measurements, reporter calibration experiments, substrate titration datasets, and target titration datasets. These files provide templates for the required spreadsheet formatting and allow users to reproduce the example analyses described in this protocol. The notebook is written for execution in Google Colab using the default Python runtime and standard scientific Python packages preinstalled in Colab, including NumPy, pandas, matplotlib, SciPy, and ipywidgets. The scripts include browser-based file-upload prompts and interactive plotting utilities to facilitate analysis by users with minimal programming experience. Additional instructions regarding input formatting, execution order, variable definitions, troubleshooting, and local execution are provided in the accompanying user manual.
